# A new protocol for speed vernalisation of winter cereals

**DOI:** 10.1101/2021.12.01.470717

**Authors:** Jin-Kyung Cha, Kathryn O’Connor, Samir Alahmad, Jong-Hee Lee, Eric Dinglasan, Hyeonjin Park, So-Myeong Lee, Dominique Hirsz, Soon-Wook Kwon, Youngho Kwon, Kyeong-Min Kim, Jong-Min Ko, Lee T. Hickey, Dongjin Shin, Laura E. Dixon

## Abstract

There are many challenges facing the development of high-yielding, nutritious crops for future environments. One limiting factor is generation time, which prolongs research and plant breeding timelines. Recent advances in speed breeding protocols have dramatically reduced generation time for many short-day and long-day species by optimising light and temperature conditions during plant growth. However, winter crops with a vernalisation requirement still require up to 6–10 weeks in low-temperature conditions before transition to reproductive development. Here, we tested a suite of environmental conditions and protocols to investigate if vernalisation can be satisfied more efficiently. We identified a vernalisation method consisting of exposing seeds at the soil surface to an extended photoperiod of 22 h day:2 h night at 10°C with transfer to speed breeding conditions that dramatically reduces generation time in both winter wheat (*Triticum aestivum*) and winter barley (*Hordeum vulgare*). Implementation of this protocol achieved up to five generations per year for winter wheat or barley, instead of the two typically obtained under standard vernalisation and plant growth conditions. The protocol has great potential to enhance training and to accelerate research, pre-breeding, and breeding outcomes focussed on winter crop improvement.

## Main

Vernalisation is the requirement for a prolonged period of cold before certain plants can transition from vegetative to reproductive development. Vernalisation thus coordinates a plant’s development with its environment. In agriculture, vernalisation maximises the growing season of a crop by enabling autumn sowing without the risk of plants transitioning to reproductive development and becoming damaged by winter conditions^1^. The longer growth duration improves crop productivity and is a common feature of many wheat (*Triticum aestivum*) and barley (*Hordeum vulgare*) cultivars grown throughout the high-yielding regions of Northern Europe, the United States, Asia, and the South Pacific. However, retaining vernalisation in elite germplasm comes at a cost for generation turnover in breeding programmes. To date, the agronomic and academic standard cereal vernalisation protocol entails 6–10 weeks at a low temperature of 2–6°C^1^. Here, we optimised the environmental conditions during vernalisation and obtained a substantial reduction in the generation time for winter wheat and winter barley. This new soil-surface protocol uses an extended photoperiod (22 h light:2 h dark) and warmer temperatures (8–10°C) than typical vernalisation treatments and was effective across a genetically diverse set of germplasm. This ‘speed vernalisation’ protocol will accelerate the pace of research, training, and pre-breeding and breeding outcomes for winter cereal crops.

Due to vernalisation, crop improvement programmes that focus on developing winter cultivars do not fully benefit from recent advances made via speed breeding, in which the photoperiod is extended to a 22-h daylength and plants are grown at 22°C during this period (Supplementary Table 1)^2,3^. Vernalisation is considered a winter response, and artificial vernalisation conditions reflect that with a standard protocol of 6–10 weeks at 2–6°C under a short-day, 8-h-light: 16-h-dark, photoperiod^1,4^. However, recent and historic evidence showed that vernalisation proceeds sometimes more efficiently at slightly warmer conditions^4–6^, which suggested to us that current vernalisation conditions may be sub-optimal. To investigate this possibility, we tested vernalisation efficiency at warmer temperatures and under short-day photoperiods (Extended Data Fig. 1a-f). As the vernalisation response is quantitative (i.e., up to the point of vernalization saturation, increasing amounts of vernalisation will lead to an acceleration in flowering time), the rate and point of completion of vernalisation can be assessed by transferring vernalising plants to floral inductive conditions and scoring flowering time. To challenge the protocols, we tested wheat cultivars with weak (e.g., cv. Claire, one copy of *VERNALIZATION1 (VRN-A1)*), moderate (e.g., cv. Buster, two *VRN-A1* copies), and strong (e.g., cv. Hereward and Charger, three *VRN-A1* copies) vernalisation requirements^7^. For all cultivars tested, vernalisation completed efficiently following 6 weeks at 10°C or 14°C (Extended Data Fig. 1a-b), with warmer temperatures leading to the development of large vegetative meristems (Extended Data Fig. 1c-f). Our results support evidence from *Arabidopsis thaliana* in which vernalisation completes in the autumn^8^, suggesting that artificial vernalisation conditions using low temperatures of 4–6°C may not be necessary. Here, raising the vernalisation temperature to 10°C for 6 weeks met all vernalisation needs of all tested cultivars, even those with long vernalisation requirements. Importantly, these new conditions of 10°C under an 8-h-light:16-h-dark photoperiod, which we refer to as warm regular vernalisation (wRV), can be supported by most controlled growth chambers.

Building on this, we were curious if there were opportunities for further optimisation. Short-day photoperiods are typically used for artificial vernalisation of cereals as they repress the expression of the long-day-activated flowering repressor *VRN2*^9^. However, there is increasing evidence that pre-vernalisation repression of flowering in cereals is a multigenic response^10,11^, prompting us to hypothesise that *VRN2* may not be a limiting factor. To test this possibility, we vernalised plants at 10°C under a 22-h-light:2-h-dark photoperiod (speed vernalisation: SV) before transfer to speed breeding (SB) conditions^2^ (SB: 22 h light:2 h dark, 22°C:17°C cycles or constant 22°C) (Fig. 1a). As with wRV, meristems remained vegetative during SV, and the plants went on to produce fully developed, fertile spikes (Extended Data Fig. 1g-n). To challenge our protocol, we used genotypes with a range of pedigrees, including Korean, European, and American wheat cultivars. We observed a reduction in generation time for many cultivars subjected to SV compared to wRV also transferred to SB (Fig. 1: Supplementary Table 2) and a substantial reduction in generation time when compared to wRV followed by regular glasshouse conditions (RG 20°C; 16 h light:8 h dark)(Extended Data Fig. 1). For example, following 4 weeks of SV, the generation time for cv. Claire was accelerated by at least 10 weeks compared to standard practice (Fig. 1 and Extended Data Fig 1a). For cv. Charger, SV enabled an acceleration of flowering following 4 weeks of vernalization from non-flowering under wRV to 125 days under SV, both transferring into SB (Fig. 1c). However, this acceleration still meant that cv. Charger generation time was longer than a cultivar with a lower vernalisation requirement (e.g., cv. Claire). The variability in the response suggested that *VRN-A1* copy number may be important, so we were interested in identifying ways to improve our protocol to mitigate this.

**Figure 1.**
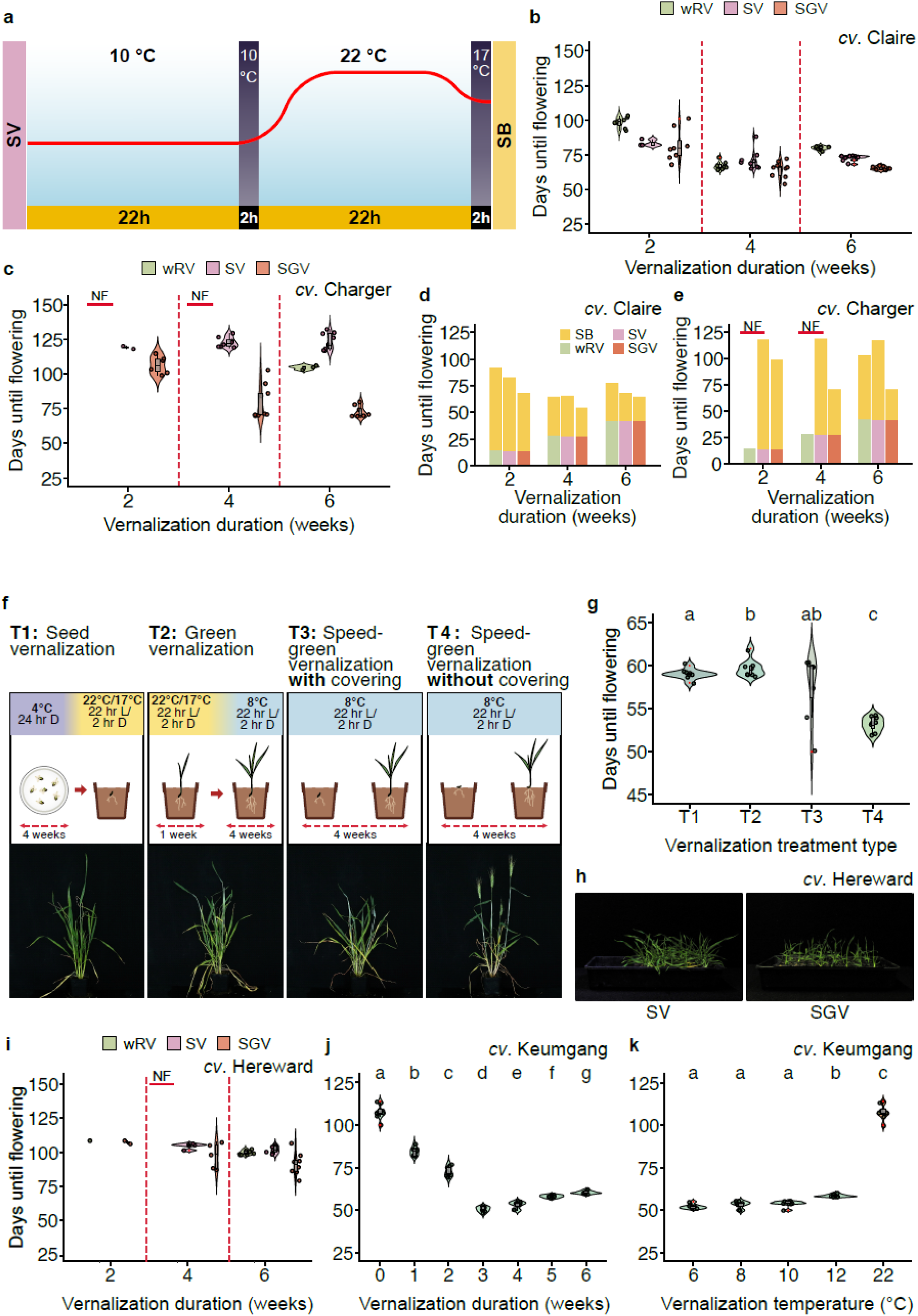
Identifying speed vernalisation conditions for wheat (Triticum aestivum) Speed (green) vernalisation (SV (SGV)) combined with speed breeding (SB) accelerates winter wheat life cycles. **a**. Scheme of SV–SB conditions. Comparison of flowering time for **b**. cv. Claire and **c**. cv. Charger between wRV, SV, and SGV treatments and transfer to SB conditions following the vernalisation duration indicated. Total time to flowering is shown (including vernalisation duration); *n* = at least 6. Fastest generation time following wRV, SV, and SGV indicated by the time the first plant flowered following transfer to SB for **d**. cv. Claire and **e**. cv. Charger. **f**. Comparison of seed treatments with example images taken when the first plant flowered under T4 conditions for the winter wheat cv. Keumgang. **g**. Flowering time following the four seed treatments (T1-4 in f) in cv. Keumgang; *n* = 7. **h**. Comparison of plant development between SV and SGV. **i**. Comparison of flowering time for cv. Hereward between wRV and SGV; *n* = at least 6. Flowering times for different durations **j**. and temperatures **k**. for cv. Keumgang under SGV. Significance is shown according to Student’s *t*-test *P* < 0.05. SV (speed vernalisation), SGV (speed-green vernalisation), wRV (warm regular vernalisation), SB (speed breeding), and vernalisation duration are included in days to flowering.

We thus used other parameters to increase the universal nature of the vernalisation protocol. Under standard practice, plants are transferred to vernalisation following 1–2 weeks of growth under regular glasshouse conditions (RG 20°C; 16 h light:8 h dark) to allow seedling establishment. Our SV protocol uses germination and growth under 22-h-light:2-h-dark photoperiods, so seedlings grow faster than with wRV. Therefore, to limit the extent of initial seedling growth, we tested if seedlings responded efficiently without pre-growth in the glasshouse. Accordingly, we subjected seeds to four treatments: T1) germination and growth at 4°C in the dark, T2) germination in SB before transfer to SV, T3) SV with the seed buried in soil, and T4) SV with the seed on the soil surface (Fig. 1f, Supplementary Table 3). Unexpectedly, vernalisation was most efficient when seeds were placed on the soil surface and exposed to light under SV conditions (Fig. 1g), which significantly reduced the time to flowering by an additional 8 days compared to the other conditions (Duncan’s multiple test, α = 0.05) for cv. Keumgang (one *VRN-A1* copy). To optimise the protocol, we tested a suite of durations and temperatures of seed-surface SV conditions using the Korean winter wheat cv. Keumgang and the American winter wheat cv. Sturdy (one *VRN-A1* copy). These experiments confirmed that 8–10°C is the most efficient and reliable vernalisation temperature (Fig. 1j-k, Supplementary Tables 4 and 5). This result revealed a seed-based aspect of the vernalisation response in cereals that is similar to dormancy in dicots^12^. The same method was also successful in cultivars with a strong vernalisation requirement (e.g., cv. Hereward and Charger) and reduced generation time by at least 4 weeks compared to SV or wRV and transfer to SB (Figure 1b-i). Importantly, the combination of seed-surface vernalisation via SV followed by SB (hereafter SGV_SB) conditions reduced the duration of vernalisation needed, with 3–4 weeks at 10°C being optimal; therefore, our protocol enables a higher throughput of plants through vernalisation compared to RV and SV.

For use in academic and industrial breeding programmes, the SGV_SB protocol should support usual plant development. Therefore, we measured multiple plant growth parameters, obtained normal seed set, and observed standard plant development, although plant height was slightly reduced (Extended Data Fig. 2). We also tested the effectiveness of the SGV_SB protocol on a wheat diversity panel regularly used in wheat breeding programmes in Korea. Of the 51 winter cultivars in the panel, 45 were fully responsive to SGV_SB conditions (Supplementary Table 6). Overall, our data demonstrate that vernalisation on the soil surface imposed by the SGV_SB protocol reduces the vernalisation requirement and generation time for many winter wheat cultivars tested and is effective even on cultivars that traditionally have a longer vernalisation requirement.

Given the similarity between the vernalisation response of wheat and barley, we hypothesised that SGV_SB may accelerate generation time in winter barley as well. Accordingly, we subjected 60 diverse winter barley cultivars originating from 34 growing regions or countries to RV_RG, RV_SB, and SGV_SB (Extended Data Fig. 3a, b). We recorded days to flowering in each experiment (Fig. 2a, b). Notably, all cultivars flowered and produced viable seeds under all conditions. Under SGV_SB, the entire population flowered substantially earlier, on average after 50 days compared to 92 days under RV_RG and 68 days under RV–SB (Fig. 2a). Therefore, in contrast to wheat, the collection of winter barley cultivars was completely responsive to the SGV_SB protocol.

**Figure 2.**
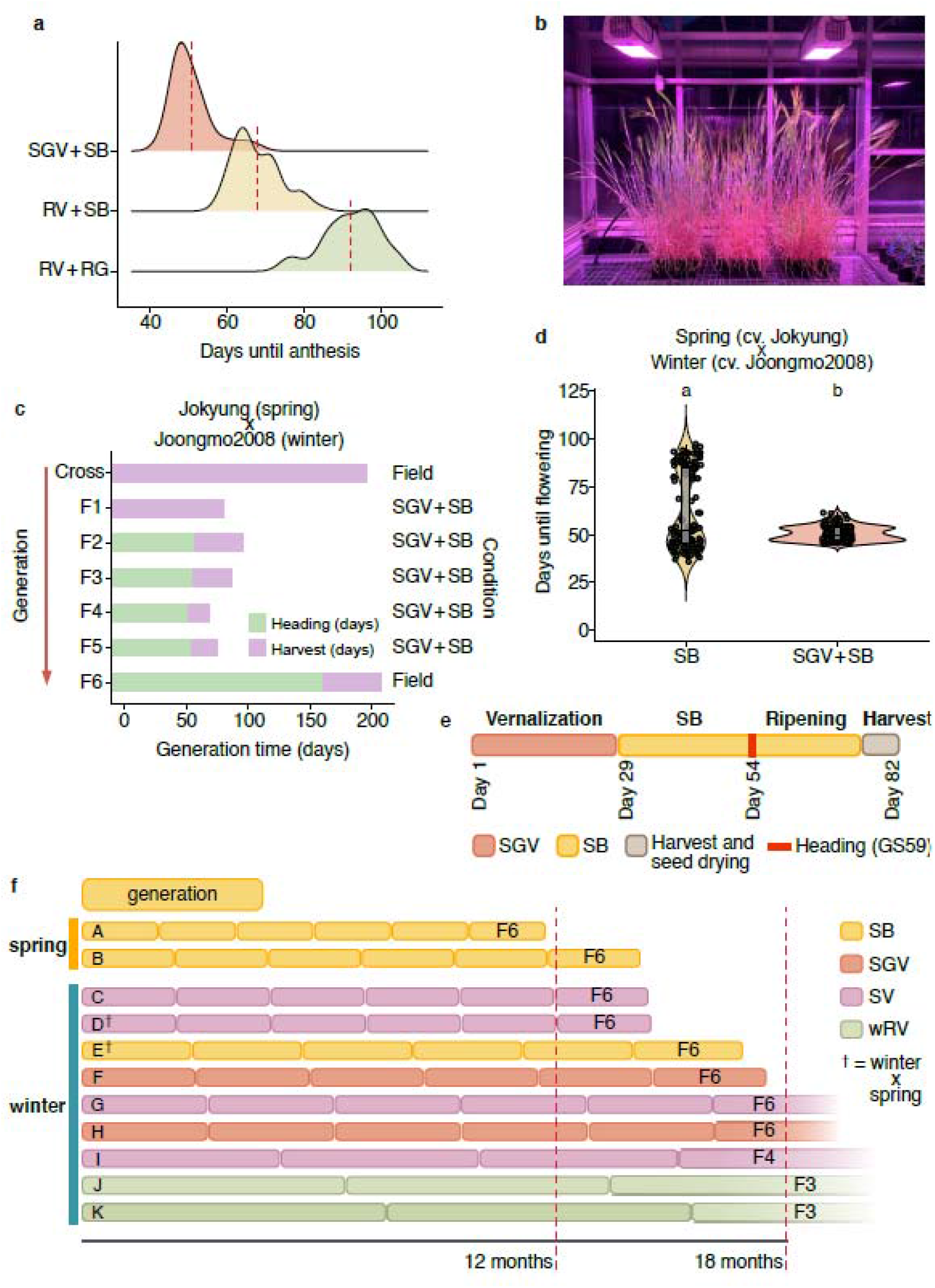
Speed vernalisation conditions for barley (Hordeum vulgare) and cereal population development. Flowering time responses of diverse winter barley accessions following various vernalization conditions. **a**. Density plots displaying days to anthesis (DTA) for 60 diverse winter barley accessions evaluated using three treatments: 1) speed vernalisation (at 8°C) and speed breeding conditions (SGV–SB), 2) regular vernalisation and speed breeding conditions (RV–SB), and 3) regular vernalisation and regular glasshouse conditions (RV–RG). **b**. Example image of barley population following SGV–SB. **c**. Population development timeline for winter x spring wheat population over six generations and **d**. flowering comparison between SB and SGV–SB for spring x winter population. **e.** An example breakdown of growth cycle under SGV–SB conditions for the same winter x spring cross in **c** and **d**. **f.** Projected generation times for different cereal types and growth conditions: A = SB Spring barley (e.g., cv. Commander, Golden Promise), B = SB Spring wheat cv. Suntop, Cadenza, C = SGV–SB Winter barley, D = SGV–SB Wheat winter x spring (using data from d), E = SB wheat winter x spring (using data from d), F = SGV–SB winter wheat cv. Claire, G = SV–SB winter wheat cv. Claire, H = SGV–SB winter wheat cv. Charger, I = SV–SB winter wheat cv. Charger, J = wRV–RG winter wheat cv. Charger, and K = wRV–RG cv. Claire. Spring generation times calculated from published data^2^. SV (speed vernalisation), SGV (speed-green vernalisation), wRV (warm regular vernalisation), SB (speed breeding).

Crossing and the subsequent development of genetically stable or inbred lines are routine practices in both breeding and research programmes. However, these techniques are particularly time consuming for populations derived from winter × spring and winter × winter crosses, as each generation must be vernalised to avoid unintended selection against the vernalisation mechanism. To test the SGV_SB protocol for use in population development, we crossed cv. Jokyung (spring) and cv. Joongmo2008 (winter) wheat cultivars in the field in Korea during the 2018–2019 winter season. Throughout the subsequent 15 months, we reached the F5 generation, with each generation taking an average of 82.4 days (Fig. 2c, Supplementary Table 7, 8). The SGV_SB protocol considerably reduced the variation in days to flowering for populations derived from winter × spring and winter × winter crosses (Fig. 2d; Extended Data Fig. 3, Supplementary Table 9). Early and more synchronous flowering across segregating or diverse germplasm can facilitate more efficient crossing and rapid generation times. We harvested at least 25 seeds from each F5 plant, which is sufficient to bulk seed in the field and subsequent evaluation. Using a projected timeline for plant generations (Fig. 2e-f), we highlight the opportunity to increase the number of plant generations within a 12–18-month period for both spring and winter wheat and barley. Impressively, the SGV_SB protocol applied to some winter cultivars reached generation turnover times similar to those of spring cultivars. Furthermore, the seed surface vernalisation is extremely high-throughput, with a density of up to 1,709 seeds per m^2^ being treated when utilising 128-cell seed trays.

Our optimised vernalisation conditions (SV and SGV) that lead to a reduction in overall generation time challenge our current understanding of the vernalisation mechanism itself, where short days reduce *VRN2* expression and cold temperatures activate *VRN1* expression^4^. Therefore, we investigated how vernalisation genes responded during SV_SB and if they differed between the seed surface (SGV) versus buried seed (SV)protocol (Fig. 3). The expression patterns of vernalisation genes followed the expected profiles in SV_SB buried seed, using cv. Claire, with *VRN1* and *VRN3* (also named *FLOWERING LOCUS T-like1* (*FT1*)) transcript levels increasing during vernalisation and *VRN2* (a locus comprised of *ZCCT1* [for *Zinc domain and CONSTANS, CONSTANS-LIKE, TOC1*] and *ZCCT2*) transcript levels decreasing (Fig. 3a-d). Notably, *VRN1* and *VRN3* expression was lower under SV compared to wRV conditions (Fig. 3a-b), indicating that these genes may not represent the exclusive route promoting flowering under SV. We observed a sustained repression of *ZCCT1* expression during and following SV when plants were transferred to SB, which was unexpected given the extended photoperiod of SV, which would be anticipated to promote *VRN2* expression (Fig. 3d). However, comparing the SV_SB conditions in cv. Hereward, which vernalised more rapidly with the seed-surface protocol, *ZCCT1* steadily increased in expression, opposite to that in the buried seed condition (Fig. 3j-k). This result indicated that considering the *VRN2* locus as a single gene is potentially misleading for our understanding of the vernalisation response and that an additional or alternative vernalisation response is activated during SGV. Further examination of this response may identify new breeding targets in the regulation of vernalisation and flowering time in cereals.

**Figure 3.**
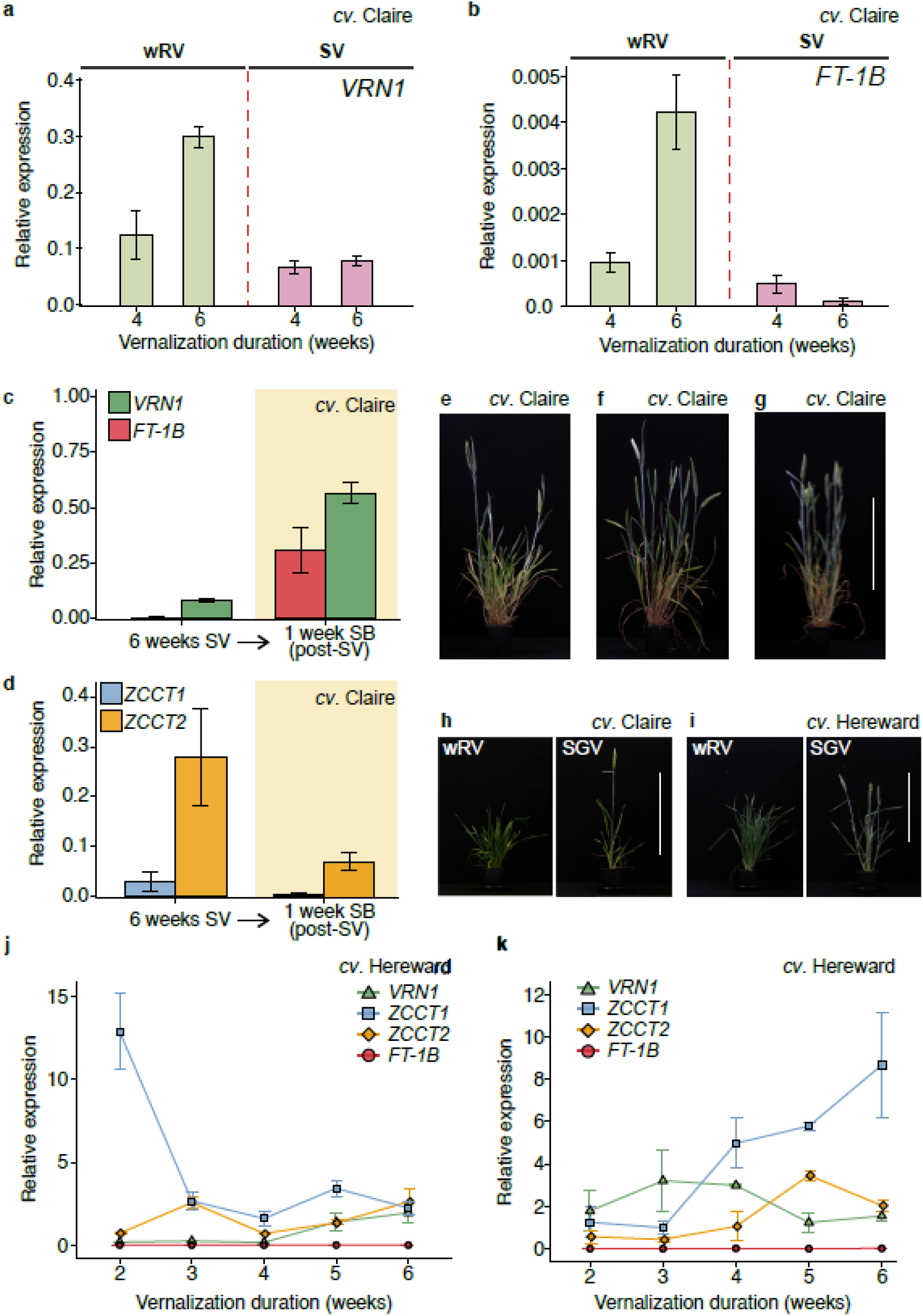
Vernalisation-related gene expression under SV and SGV conditions. In cv. Claire, **a**. Comparison of expression of *VRN1* between wRV (green) and SV (pink) conditions. **b**. Comparison of expression of *FT1-B* between wRV (green) and SV (pink) conditions. **c**. Expression of *VRN1* (green) and *FT1-B* (red) between 6 weeks of SV and then 6 weeks of SV and 1 week in SB. **d**. Expression of *ZCCT1* and *ZCCT2* between 6 weeks of SV and then 6 weeks of SV and 1 week in SB. Representative images following different vernalisation treatments. For cv. Claire following SV_SB with 3 weeks **e**., 4 weeks **f**., and 5 weeks **g**. vernalisation treatment. Comparison between wRV and SGV conditions for cv. Claire following 4 weeks vernalization and photographed at the same age **h**. and cv. Hereward following 6 weeks vernalization and photographed at the same age **i**. All conditions result in vernalised plants. Scale bar is 30 cm. In cv. Hereward, expression of *VRN1, ZCCT1, ZCCT2*, and *FT-1B* for **j**. SV and **k**. SGV. All show three biological replicates that each comprise at least three plants, with standard error of the mean. SGV (speed-green vernalisation), wRV (warm regular vernalisation), duration in weeks refers to vernalisation duration experienced.

The time required for the introgression and stacking of novel alleles remains a bottleneck in the development of improved cultivars. One of the limiting factors is the vernalisation requirement, which imposes a biological constraint on generation time. Here, we identify a method to reduce generation time for winter wheat and barley germplasm using alternative vernalisation conditions. The protocols we developed identify the environmental parameters that can be further modified to account for local or genotypic variation in vernalisation efficiencies and therefore offer a framework to universally reduce generation times in winter cereals. The approach could also be adapted to other winter crops (e.g., canola [*Brassica napus*]) or vegetables with a vernalisation requirement to reduce generation time and accelerate breeding outcomes.

## Materials and methods

### Cultivars used

Wheat and barley cultivars used in this study are in Supplementary Table 10.

#### European winter wheat assessment and gene expression (assessed at The University of Leeds, UK)

##### Conditions used: Wheat (*Triticum aestivum*)

*wRV_RG:* 8 h light:16 h dark 10°C into 16 h light:8 h dark 22°C

*wRV_SB:* 8 h light:16 h dark 10°C into 22 h light:2 h dark 22°C

*SV (and SGV) _SB:* 22 h light:2 h dark 10°C into 22 h light:2 h dark 22°C

Please note warmer conditions were used in wRV than are classically used in RV to enable direct photoperiod comparison for phenotype and gene expression analysis.

Seeds were germinated for 2 days in darkness at 4°C in 9-cm petri dishes that had a layer of filter paper saturated with 5 mL dH_2_O. The germinated seeds were transferred to 3 × 3-cm cell pots of JIC cereal mix^4^ and placed under vernalisation conditions in Snijders MICROCLIMA MC1000 cabinets. Plants were watered when required, and no additional nutrients were added. At 1- or 2-week intervals, plants were sampled for gene expression and apex analysis (see below), and three plants were transferred to glasshouse growth conditions for RG (PhytoLux Plessey; model ATTIS-7) and SB (Heliospectra; model MITRA). Plants were placed in cereal mix (as above) in 9 × 9-cm pots. Plants were sampled 1 and 2 weeks after transfer to SB conditions. Flowering time was recorded as half-ear emergence (Zadok scale 55). Plants that did not flower after 170 days were recorded as “non-flowering.” A control group using the same four genotypes (*n* = 10, *s* = 40) was grown under constant SB conditions, and a representative for each genotype was imaged once the plant had reached maturity.

### Speed-green vernalisation

Seeds were placed in 9-cm petri dishes containing filter paper and 5 mL dH_2_O and kept in complete darkness at 4°C for 48 hours. Seeds were then placed on the soil surface (John Innes cereal mix) in a P24 seedling tray. Care was taken to press the seed into the soil surface while ensuring the seed remained uncovered and exposed to the light. Plant trays were watered from the base, and a spray bottle of dH_2_O was used to mist the soil surface; particular care was taken misting the soil during the first week of growth, when the roots were anchoring into the soil. At set weekly intervals, plants were moved into SB conditions and flowering was recorded.

### Gene expression samples

Leaf samples from three plants per biological replicate and for three biological replicates (*n* = 3) were taken at each sampling stage. Samples were taken 1 h after lights on. To investigate gene expression during SV, leaf tissue was sampled at 1, 2, 3, 4, 5, 6, 7, and 8 weeks of growth under SV conditions. Total RNA was extracted using the Spectrum^™^ Plant Total RNA Kit (Sigma-Aldrich) following the manufacturer’s recommended protocol. RNA samples were treated using RQ1 RNase-Free Dnase (Promega), and first-strand cDNA synthesis primed with Oligo dT was processed using SuperScript^™^ III Reverse Transcriptase (Invitrogen) and RNaseOUT^™^ (Invitrogen). The cDNA was diluted (1:10), and quantitative reverse transcription PCR (RT-qPCR) was performed using the CFX96^™^ Thermal Cycler (Bio-Rad) and GoTaq^®^ Master Mix (Promega). Primers used are provided in Supplementary Table 11. Expression levels of the genes of interest were calculated relative to TraesCS5A02G015600 following the 2^^−ΔCT^ format (where ΔCT =*GOI* CT – TraesCS5A02G015600 CT).

Apex samples: three plants were dissected for each apex sample (*n* = 3) and imaged on a Keyence microscope, and apex length was measured using ImageJ^13^.

#### Methods for Wheat (*Triticum aestivum*) (assessed at RDA, S. Korea)

##### Conditions used: Wheat (*Triticum aestivum*)

*wRV–SB:* 8 h light:16 h dark 10°C into 22 h light:2 h dark 22°C:17°C

SGV_RG: 22 h light:2 h dark 10°C into 16 h light:8 h dark 22°C:17°C

*SGV–SB:* 22 h light:2-h dark 10°C into 22 h light:2 h dark 22°C:17°C

### Optimization of vernalization treatment method

Seeds were germinated in 9-cm petri dishes containing 10 mL of water and at 4°C for 3 days under dark conditions. Then, seeds were transferred to 25°C until they reached the growth stage 07 and about 0.2-mm coleoptile length^14^. For all experiments, each square pot (7(L) × 7(W) × (7H)) used was filled with 245 mL of soil, which is a mixture of paddy rice soil (Punong, Korea) and horticulture soil (Seoul-Bio, Korea) at a 2:1 ratio. To find an optimum vernalisation treatment method, four cold treatment methods were applied using the Korean winter variety Keumgang as follows: T1) seed vernalisation, T2) green vernalisation, T3) speed-green vernalisation with covering soil, and T4) speed-green vernalisation without covering soil about 1.5 cm in depth. Regarding the seed vernalisation method (T1), germinated seeds were placed in a 4°C refrigerator for 4 weeks without light and cultivated under speed breeding (SB) conditions. For green vernalisation (T2), plants initially placed under SB for 1 week were further grown under 22-h-light:2-h-dark photoperiod cycles at 8°C for 4 weeks. Concerning the speed-green vernalisation with (T3) and without (T4) covering soil, germinated seeds were sown and grown at 8°C for 4 weeks under 22-h-light:2-h-dark cycles. LED lights (red 8: blue 3: white 2, Estech LED, Korea, Extended Data Fig. 2b-c) in SGV condition were set to 410 μmol m^−2^ s^−1^ of illumination. All plants were then transferred to SB (2,500 μmol m^−2^ s^−1^ of light) to evaluate the time to flowering after vernalisation treatment; *n* = 8 plants for each evaluation. Spectral measurement of light composition was performed using the MK350N handheld spectrometer (UPRtek, Miaoli county, Taiwan).

### Optimization of vernalisation temperature and periods

To optimize the cold treatment protocol for speed vernalisation, temperatures ranging from 6–12°C were applied for 2–6 weeks under SGV without covering soil (SGV condition 4). Six plants were then transferred to SB conditions to examine days from germination to heading. The growth stages (GS32 and GS59) were recorded as reported by^14^, and plant height was measured from the ground to the start point of the spike.

### Evaluation of heading time of genetic resources

To investigate days from germination to heading time of wheat genetic resources, germinated seeds were placed in the cell of a 72-cell tray (34 mL/cell) and cultivated under the speed-green vernalisation without covering soil condition (SGV condition 4). Plants were grown under SGV for 4 weeks, followed by transfer to SB and observation until heading time (GS59), *n*= 4.

### Plant density

SGV was conducted under a range of planting densities and supported very-high-throughput vernalisation. When testing seed trays, the 72-cell seed tray, providing a total number was 466 plant/m^2^, and 105-cell tray (680 plant/m^2^) both supported normal plant growth and development.

### Development of breeding materials

To evaluate actual application of SGV in a breeding programme, 10 F1 seeds were obtained from a cross derived between cv. Jokyung (spring) and cv. Joongmo2008 (winter) wheat varieties in the field in the 2018–2019 winter season. Germinated seeds from the F1 to F5 generation were placed in the cell of a 72-cell tray (34 mL/cell) and cultivated under the speed-green vernalisation without covering soil condition (SGV condition 4). Plants were grown under SGV for 4 weeks, followed by transfer to SB until harvest. After harvest, seeds were dried in the glasshouse for 7 days for the next generation. For F6 generation, 25 seeds of each line were planted in the field at the National Institute of Crop Science, Miryang Korea (35.3° N; 128.5° E).

### Barley collection (assessed at The University of Queensland, Australia)

#### Conditions used: Barley (*Hordeum vulgare*)

*RV–RG:* 8 h light:16 h dark 6°C into 16 h light:8 h dark 22°C

*RV–SB:* 8 h light:16 h dark 6°C into 22 h light:2 h dark 22°C:17°C

*SGV–SB:* 22 h light:2 h dark 8°C into 22 h light:2 h dark 22°C:17°C

### Experimental design and treatments

The barley accessions were evaluated in three experiments: 1) regular vernalisation and regular glasshouse (RV–RG), 2) regular vernalisation and speed breeding (RV–SB), and 3) speed vernalisation and speed breeding (SGV–SB). Under regular vernalisation conditions, plants received a standard vernalisation treatment at 6°C for 6 weeks under 8-h-light: 16-h-dark photoperiod cycles. For speed vernalisation (SGV), plants were exposed to 8°C for 4 weeks under 22-h-light:2-h-dark cycles. Vernalisation was performed in a fully enclosed walk-in growth cabinet fitted with LED growth lights (Heliospectra, model E602G). For the regular vernalisation (RV), seeds were sown directly into 100-cell trays (18 mL per cell), covered with UQ23 potting mix^15^, watered, and moved into the vernalisation chamber. For speed vernalisation (SGV), seeds were pre-germinated in petri plates and then placed onto the surface of the potting mix. Three seeds per accession were transplanted into a single cell of the tray for vernalisation. To retain moisture in the cells during vernalisation, the 100-cell trays were placed inside a sealed bottom tray with two to three small drainage holes. Trays were lightly watered daily during the vernalisation process.

After vernalisation, the bottom trays were removed and filled with UQ23 potting mix and Osmocote^®^ slow release fertiliser (at a rate of 2 g per litre) to provide developing plants with sufficient media and resources. For the regular glasshouse conditions, plants were grown in a temperature-controlled glasshouse (22/17°C, light/dark) under a natural 12-h diurnal photoperiod. For the speed breeding treatment, trays of barley plants were moved to a temperature-controlled glasshouse (22/17°C, light/dark) fitted with Heliospectra LEDs using a 22-h-light:2-h-dark photoperiod^15^. The day of anthesis for each accession was recorded as the first spike to reach awn-peep stage (GS49).

### Benchmarking diversity of the 60 winter barley accessions

A diverse panel of barley accessions acquired from Australian Grains Genebank Collection (AGG, *n* = 806) was genotyped using the Illumina Infinium 40K XT SNP chip assay (InterGrain and AgriBio-Victoria) and returned a total of 12,599 SNP markers. Polymorphic markers with known chromosome positions were used to investigate the genetic diversity of the barley accessions (9,221 high-quality SNP markers with <10% missing data and <10% heterozygosity; and 737 barley lines with <10% missing values). To investigate the population structure, we calculated the pairwise Roger’s distances between the accessions using ‘SelectionTools’ (downloadable at http://population-genetics.uni-giessen.de/~software/) implemented in R. Principal coordinate analysis based on the Roger’s genetic distance matrix and k-means clustering was performed and plotted using ggplot2^16^ in R.

### Statistical analyses

Variation in DTA for the barley accessions in each vernalisation treatment was visualised in the form of density plots, generated using ggjoy package in R. To determine if DTA differed across the three treatments, one-way analysis of variance (ANOVA) was performed. Tukey’s multiple comparison test (Tukey’s HSD test) was then performed to evaluate the effect of each treatment on DTA. HSD test applies appropriate adjustments to the mean for each treatment suitable to multiple testing^17^. The analysis was performed using (Agricolae package) in R. To investigate the relationship between barley flowering behaviour across vernalisation treatments, the Pearson’s correlation coefficient (*r*) was calculated for DTA. The degree of correlation was also tested for significance (*P*-value; *α* = 0.05). The analysis was performed using corrgram and corrplot packages in R.

## Acknowledgements

This research was supported by the Research Program for Agricultural Science and Technology Development (Project No. PJ011202) Rural Development Administration. L.T.H received funding from the Australian Research Council (ARC), project codes DP190102185 and LP170100317. Genotyping of the winter barley accessions at The University of Queensland was funded through the Grains Research and Development Corporation (GRDC), project code UOQ2005-012RTX. S.A was supported a GRDC Postdoctoral Fellowship, project code UOQ1903-007RTX. L. E. D. at the University of Leeds received funding from UKRI FLF MR/S031677/1 and the Rank Prize Funds New Lecturer Award.

## Author contributions

L.T.H., D.S. and L.E.D. conceived and supervised the project. J.H.L., L.T.H., D.S. and L.E.D. designed the experiments. J.K.C. K.O.C., D.H. and K.M.K. investigated flowering time of wheat. S.A. and E.D. investigated flowering time of barley. J.K.C, H.P., S.M.L., Y.K. and J.M.K. developed wheat breeding materials. J.K.C., K.O.C. and S.W.K. analysed the data. L.T.H., D.S. and L.E.D. wrote the manuscript. All the authors discussed the results and contributed to the manuscript.

---

**Extended data figure 1:**
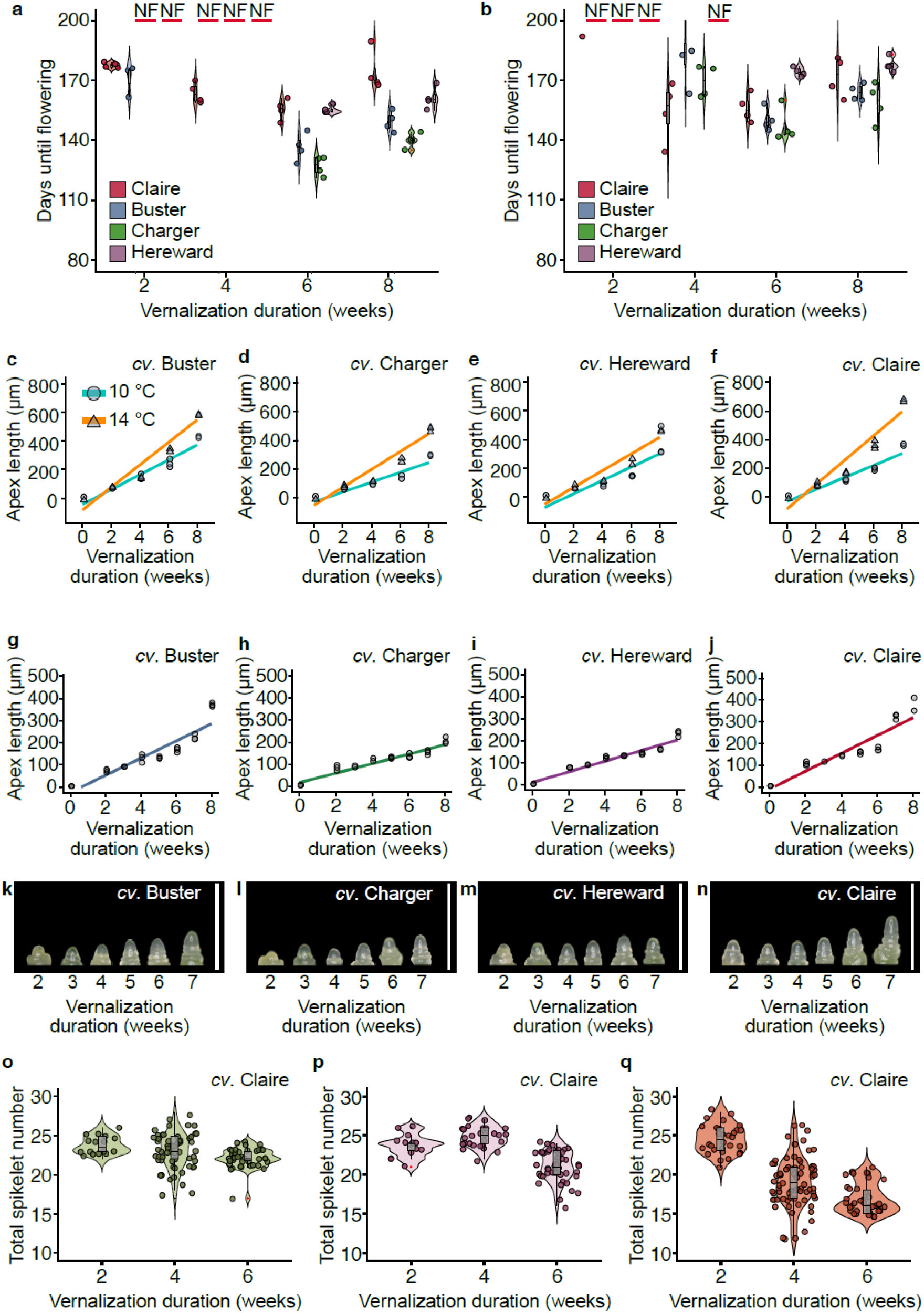
Plant development under different vernalisation temperatures for four European winter wheat cultivars. Days to flowering (GS59) for winter wheat cultivars Claire, Buster, Charger, and Hereward vernalising under **a**. 10°C or **b**. 14°C short-day (8 h light:16 h dark) photoperiod and transferred at the weeks indicated on the x-axis to long-day (16 h light:8 h dark) glasshouse conditions with constant 20–22°C. Vernalisation duration is included in the flowering time. *n* = at least 3, NF = No flowering by date indicated. **c**-**f** Apex lengths from the same conditions; *n* = 3. Apex length following SV for cultivars **g**. Buster, **h**. Charger, **i**. Hereward, and **j**. Claire; *n*= 3 with representative images between weeks 2 and 7 of SV for each cultivar, scale bar represents 1000 μm, for cultivars **k**. Buster, **l**. Charger, **m**. Hereward, and **n**. Claire. Spikelet counts from plants moved following 2, 4, or 6 weeks of vernalisation for cv. Claire for **o**. wRV_SB, **p**. SV_SB, and **q**. SGV_SB. *n* = at least 8.

**Extended data figure 2:**
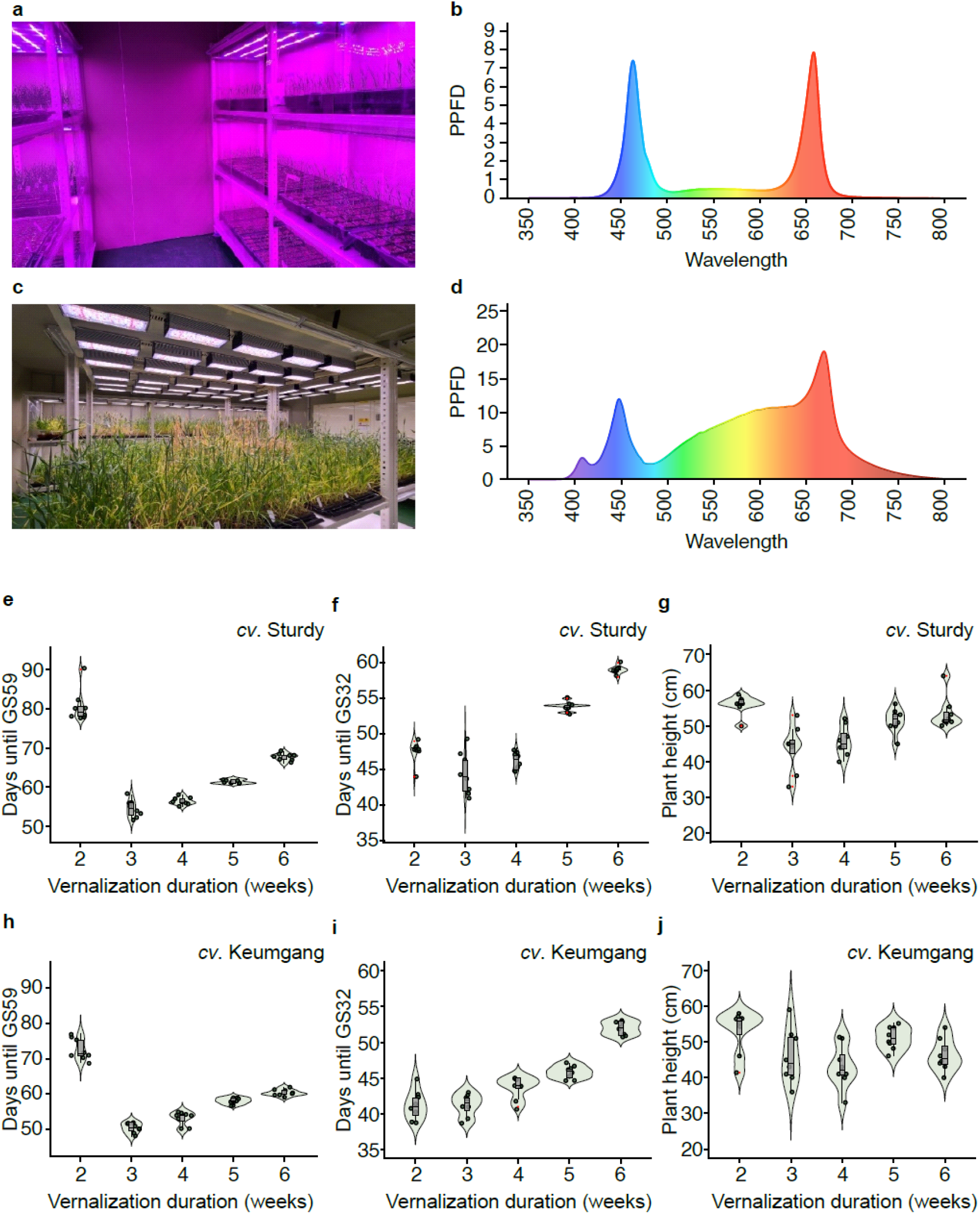
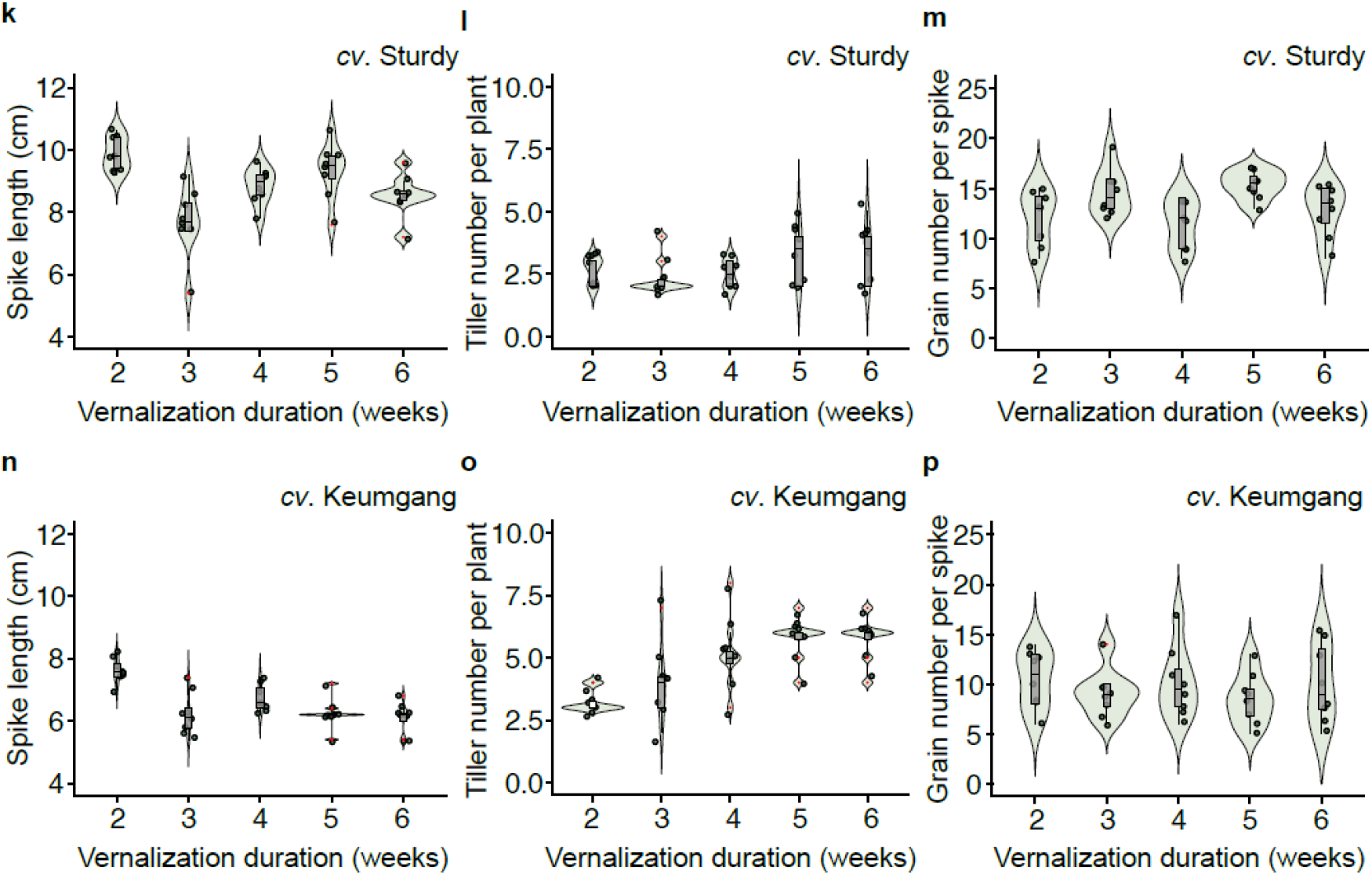
An example of SGV_SB conditions using LED lighting. **a**. Representative image of plants in SGV conditions with **b**. showing the spectral measurement of light composition in SGV. **c.**Representative image of plants in SB conditions and **d**. showing the spectral measurement of light composition SB. Spectral measurements were taken using the RS-3500 spectroradiometer from spectral Evolution Inc. X-axis values are wavelengths in nanometres, and y-axis represents proportion (1 unit = 0.1 proportion). Under SGV–SB conditions, wheat plants followed standard development shown over a 2- to 6-week vernalisation treatment for cv. Sturdy: **e**. Growth stage 59, **f**. days until GS32, **g**. plant height for cv. Keumgang, **h**. growth stage 59, **i**. days until GS32, **j**. plant height for cv. Sturdy, **k.** Spike length, **l**. tiller number per plant, and **m**. grain number per spike; and for cv. Keumgang: **n**. spike length, **o**. tiller number per plant, and **p**. grain number per spike, where *n* = 6.

**Extended data figure 3:**
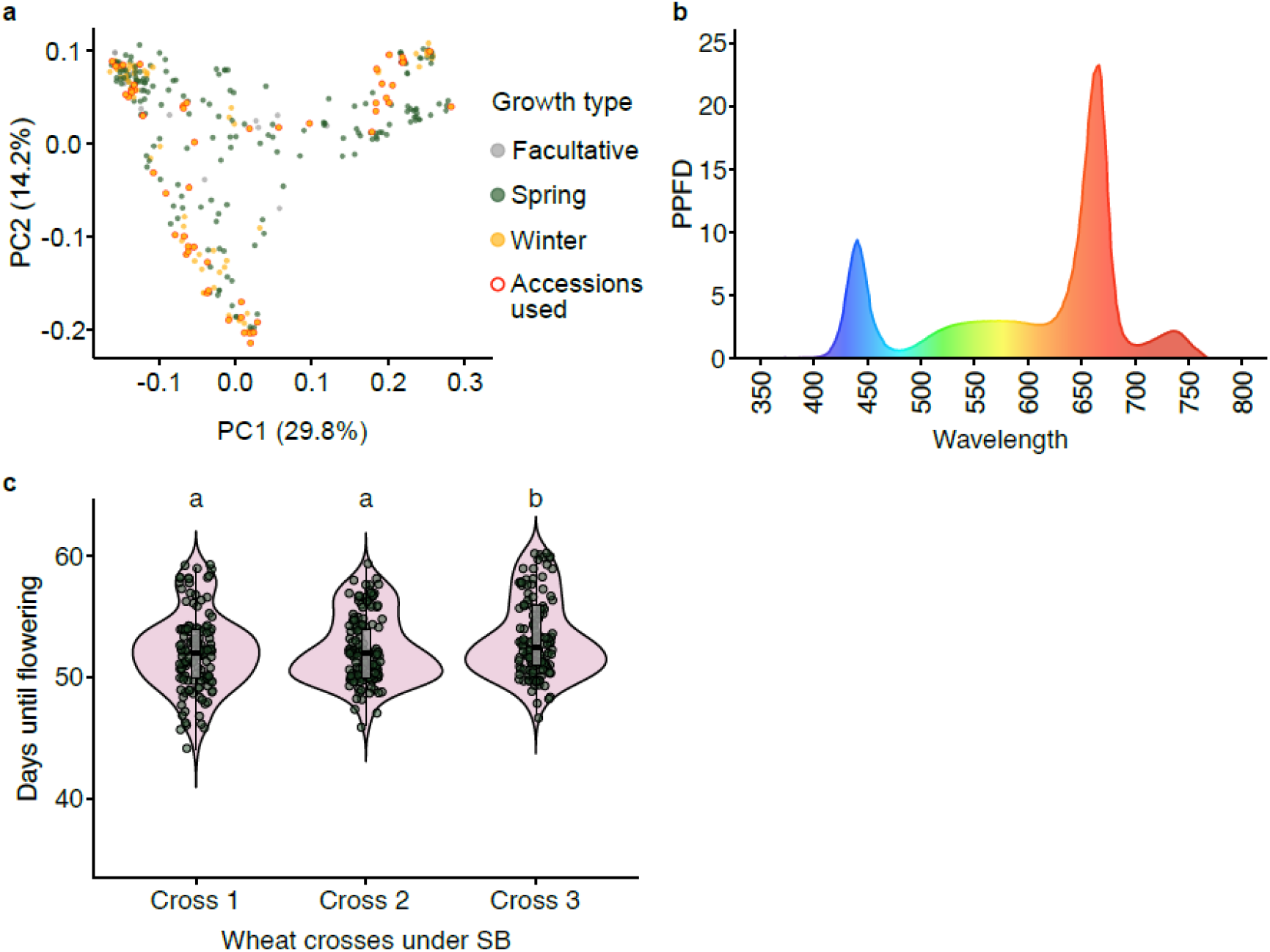
Accelerated generation times also achieved for barley and breeding populations. **a**. Principal component analysis (PCA) of barley cultivars used in Figure 2. **b**. Spectral measurement of light composition during SGV for barley and **c**. flowering distribution for three additional spring x winter or winter x winter populations following SGV–SB; crosses 1 and 2 are spring x winter populations, and cross 3 is a winter x winter population.

